# The effects of soleus push up on glucose tolerance among individuals with prediabetes

**DOI:** 10.1101/2024.11.14.623602

**Authors:** Dávid Elek, Miklós Tóth, Balázs Sonkodi, Pongrác Ács, Gábor L. Kovács, Csaba Melczer

## Abstract

The objective of this study was to assess the efficacy of a previously invented soleus-dominant exercise in reducing postprandial blood glucose levels in individuals with prediabetes and to evaluate the feasibility of incorporating this soleus push up exercise into their daily routine. It seems that performing the mostly soleus-involved activity in a sitting position during the oral glucose tolerance test results in ∼32% lower postprandial glucose excursion than the control results. This effect is also present in the absence of electromyographic feedback. Hereby we conclude that performing this repetitive prolonged contractile muscle activity may help metabolic regulation even in prediabetic people without the need of a laboratory setting. Therefore soleus push up is a potent exercise method that can be effectively completed in a home setting or work environment. However, further validation of this treatment method should be studied in a larger sample size.

## INTRODUCTION

A It is becoming increasingly evident that the major technological advances of recent decades, through their impact on the social fabric and social connections, are contributing to a downward trend in the physical activity of the population in developed countries. In 2016, it was observed that 28% of the world’s adult population and 81% of adolescents were below the WHO guidelines for health promotion of 150 minutes of moderate-intensity physical activity per week for adults or 75 minutes of high-intensity physical activity per week for adults and 60 minutes of moderate-intensity physical activity per day for adolescents [1].

It is worth noting that a caloric surplus, which can result from a combination of reduced physical activity and a calorie-dense diet, may contribute to the development of obesity and a high BMI. It is estimated that more than 10% of the world’s population is overweight, which is known to be a significant risk factor for several diseases, including cardiovascular disease, orthopedic conditions and type 2 diabetes mellitus. Hungary is placed fourth in the ranking of countries for overweight in the adult population among those in the Organization for Economic Co-operation and Development (OECD). It would be fair to say that a significant proportion of the adult population in the country is overweight, which is a factor that can contribute to a range of health issues, including cardiovascular disease, cancer and metabolic disorders such as type 2 diabetes.

It is estimated that in Hungary, in addition to 110,000 undiagnosed cases of type 2 diabetes, around 661,000 people have diabetes, which represents a prevalence of 9.1%. It is also worth noting that 237,000 people in Hungary had impaired glucose tolerance (IGT) and 306,000 people had elevated fasting blood glucose (IFG). IGT and IFG are the precursors to type 2 diabetes [2]. In 2019, the social security system allocated a total of HUF 8 billion for the provision of specialized care for patients with diabetes mellitus. Of this amount, 70%, or HUF 5.8 billion, was attributed to the funding of type 2 diabetes care [3, 4].

Many studies have sought to treat prediabetes, resulting from the lifestyle attitudes mentioned above, with varying degrees of success. However, a research study published in 2022 at the University of Houston found that repeating a previously unstudied exercise method called the “Soleus Push up” (SPU) for three hours resulted in a notable improvement in VLDL and triglyceride levels and glucose tolerance in the participants. It seems that this form of exercise may have induced a large systemic effect by isolated contraction of a muscle (musculus soleus) with a predominantly slow oxidative fiber composition, which accounts for ∼1% of the total body mass. In addition, it appears that it may have enhanced local oxidative metabolism. Given that the movement can be performed in a sitting position for extended periods without fatigue, it has the potential to be a convenient yet effective method of metabolic enhancement that could be incorporated into daily activities. Upon reviewing the demographic data of the participants in the trial, it became evident that there was a considerable range in the observed parameters. Nevertheless, the findings indicated that SPU was effective in improving glucose regulation at low physical activity levels [5].

The aim of our study is to investigate the potential benefits of SPU for people with prediabetes and to analyze its clinical applications. The additional purpose of this paper is to present a pilot study to provide a basement for further work.

## RESULTS

### Characteristics of the participants

The participants (n=10) consisted of 6 male and 4 female volunteers with an average age of 53.3 ± 2.7 years.

### Performing SPU results in improved postprandial glucose tolerance

The effect of performing SPU on blood glucose levels during the two-hour long glucose tolerance test is described in Table 1. No significant difference was observed between the group means for fasting blood glucose levels. A comparison of the means of the blood glucose values measured during the two-hour oral glucose tolerance test (OGTT) during the intervention compared to those of the control OGTT revealed a significant difference from the third sampling time onwards and lasted until the last measurement time. Significant reductions in blood glucose levels were observed in both groups compared to the control. The graphical design of Table 1, Figure 1 and Figure 3 was inspired by Hamilton’s study with the aim of better comparison of the results.

**Table 1.** Blood glucose levels at each measurement point Mean ± SEM. Mixed effect model followed by Tukey’s multiple comparison test was used to evaluate the effect of soleus push up (SPU) on blood glucose values during the oral glucose tolerance test (OGTT). *p < 0.05, **p < 0.005, ***p < 0.001 versus control measurement. The difference between SPU and control is represented at each time point (Δmmol/l).

| OGTT | With EMG feedback |  |  | Without EMG feedback |  |  |
| --- | --- | --- | --- | --- | --- | --- |
| Time (minutes) | Control (n=5) | SPU (n=5) | Δmmol/l | Control (n=5) | SPU (n=5) | Δmmol/l |
| 0 | $6.2 \pm 0.1$ | $6.2 \pm 0.1$ | $-0.1 \pm 0.1$ | $6.3 \pm 0.1$ | $6.2 \pm 0.1$ | $-0.1 \pm 0.1$ |
| 15 | $8 \pm 0.2$ | $7.7 \pm 0.2$ | $-0.3 \pm 0.2$ | $7.8 \pm 0.1$ | $7.6 \pm 0.1$ | $-0.2 \pm 0.1$ |
| 30 | $10.4 \pm 0.5$ | $9.1 \pm 0.3$ | $-1.3 \pm 0.3^*$ | $10.4 \pm 0.3$ | $9.5 \pm 0.3$ | $-1 \pm 0.2^*$ |
| 45 | $11.6 \pm 0.4$ | $9.8 \pm 0.4$ | $-1.8 \pm 0.2^{***}$ | $11.4 \pm 0.4$ | $10.1 \pm 0.3$ | $-1.3 \pm 0.3^*$ |
| 60 | $11.9 \pm 0.4$ | $10 \pm 0.5$ | $-1.8 \pm 0.1^{**}$ | $12.1 \pm 0.3$ | $10.4 \pm 0.3$ | $-1.7 \pm 0.3^*$ |
| 75 | $12.2 \pm 0.4$ | $9.8 \pm 0.4$ | $-2.5 \pm 0.2^{***}$ | $11.8 \pm 0.3$ | $10.1 \pm 0.4$ | $-1.7 \pm 0.4^*$ |
| 90 | $12 \pm 0.4$ | $9.4 \pm 0.5$ | $-2.6 \pm 0.3^{**}$ | $11.4 \pm 0.3$ | $9.6 \pm 0.5$ | $-1.8 \pm 0.3^*$ |
| 105 | $11.1 \pm 0.5$ | $8.8 \pm 0.4$ | $-2.3 \pm 0.4^*$ | $10.7 \pm 0.4$ | $9.3 \pm 0.4$ | $-1.4 \pm 0.4^*$ |
| 120 | $10 \pm 0.5$ | $8.3 \pm 0.4$ | $-1.7 \pm 0.4^*$ | $10.1 \pm 0.3$ | $8.7 \pm 0.4$ | $-1.5 \pm 0.3^*$ |

**Figure 1.**
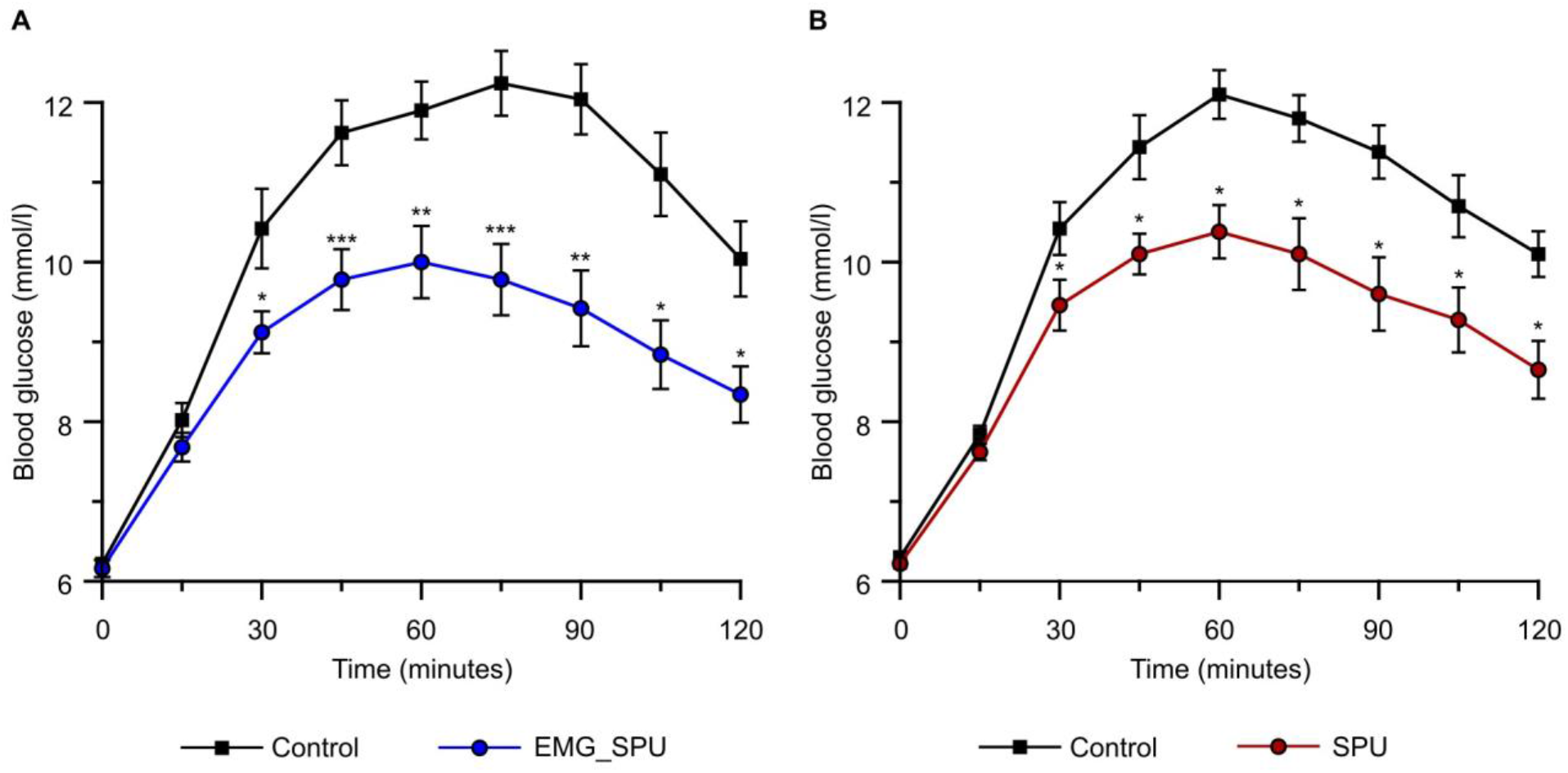
Performing soleus-dominant exercise is an effective way of immediately achieving improved glucose tolerance. Comparing the effect of SPUs on the group blood glucose level during the 120-min glucose tolerance test. (A) Performing soleus push up while continuously receiving EMG feedback about the soleus muscle activity (EMG_SPU) versus control measurement (Control) (n=5). (B) Performing soleus push up while not receiving EMG feedback (SPU) versus control measurement (Control) (n=5). Mean ± SEM. *p<0.05, **p<0.005, ***p<0.001 versus control measurement. Mixed effect model followed by Tukey’s multiple comparison test was used to evaluate the effect of SPUs. See Table 1 for detailed comparison at each time point.

The effect of SPU was observed in both groups from the 30th minute of the oral glucose tolerance test until the final measurement time point. During the period of the OGTT, the blood glucose of the group that performed the intervention with EMG feedback was, on average, 2 mmol/L lower than that of the control measurement. In comparison, the group that performed SPU without EMG feedback exhibited a mean reduction of 1.49 mmol/L during SPU compared to the control measurement. A visual comparison of the means of blood glucose values during the intervention versus control measurements for the EMG feedback-guided group is shown in Figure 1A and a comparison of the means of blood glucose values during the intervention versus control measurements for the group that did not receive continuous EMG feedback Figure 1B.

The trapezoidal rule was employed to ascertain the incremental area under the curve (iAUC), defined by the mean of the blood glucose values during the OGTT. of the control oral glucose tolerance test and the mean of the blood glucose values measured during the oral glucose tolerance test during SPU, for both the group performing SPU with EMG feedback and the group performing SPU without EMG feedback. The difference in the iAUC of the control OGTT and the intervention OGTT for the group performing SPU with EMG feedback is illustrated in Figure 2A, while the corresponding figure for the group performing SPU without EMG feedback is Figure 2B. The repetitive soleus contractions resulted in a 37% reduction in glucose iAUC of the group performing the SPUs with EMG feedback. A -26% change was observed in the group that did not use external feedback during the SPUs.

**Figure 2.**
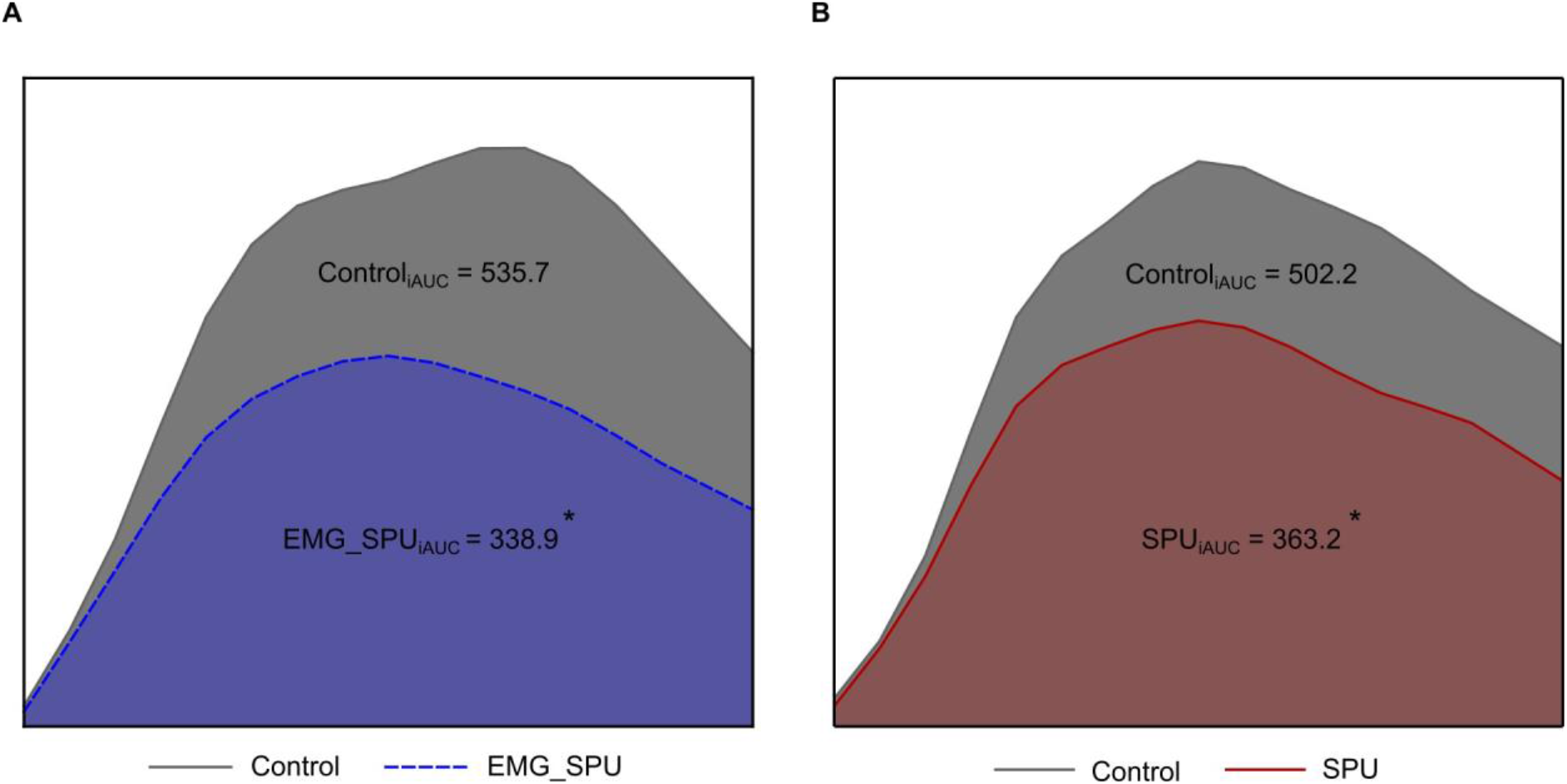
Repetitive soleus activity decreases the blood glucose incremental area under the curve by ∼32%. Data represented as incremental area under the curve (iAUC) calculated by using the trapezoid method. Coloured areas show the blood glucose iAUC change (A) in the group using continuous EMG feedback (EMG_SPU) versus control (n=5) and (B) in the group without EMG feedback (SPU) versus control (n=4). Mixed effect model followed by Tukey’s multiple comparison test was used to evaluate the difference of the iAUC between the interventional and the control measurement. *p<0.05 versus control measurement.

**Figure 3.**
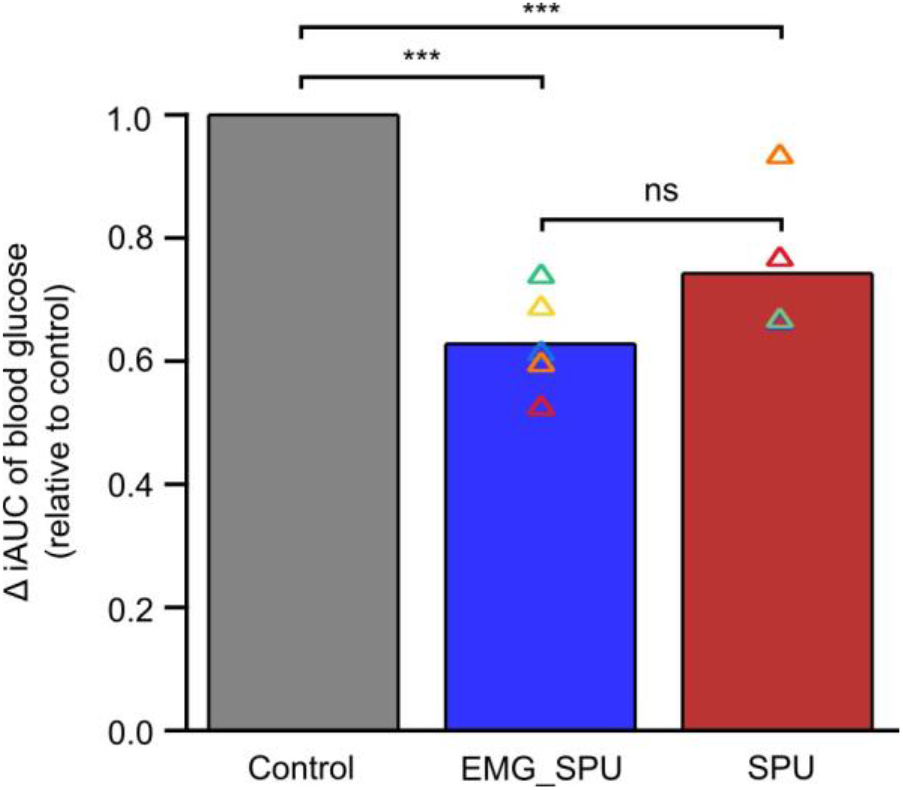
prolonged soleus muscle activity improves glucose tolerance with and without electromyographic feedback. Coloured triangles represent each individual glucose iAUC when performing soleus push ups relative to the control measurement. Bar charts summarize the mean change in the group iAUC relative to the control results in the group that used EMG feedback (EMG_SPU) and in the group that did not use external biofeedback while performing the SPUs (SPU). Mixed effect model followed by Tukey’s multiple comparison test was used to evaluate the difference of the iAUC between the interventional and the control measurement. ***p<0.001 versus control. ns=p>0.05.

It was observed in both groups that performing prolonged soleus muscle activity significantly reduced the blood glucose iAUC compared to the control results. The smallest iAUC reduction at 10% was observed in the group not using EMG feedback during the contractile activity while an individual achieved 47% reduction in the group that used continuous EMG feedback. No significant difference was observed in the extent of reduction in the area under the curve between the two groups (Figure 6).

## DISCUSSION

The objective of our study was to ascertain the impact of SPU exercise on glucose regulation in individuals with prediabetes. Additionally, we sought to examine the feasibility of integrating SPU into daily life, given that continuous EMG feedback was not available for one group.

Our study differed significantly from the one conducted by Hamilton and colleagues [5], which formed the basis for our research. While Hamilton and colleagues investigated the efficacy of SPU in healthy individuals, our study focused on examining the impact of SPU in individuals with prediabetes.

Hamilton et al. divided the participants into two groups based on their activity levels, which were defined as 1.3 and 1.7 metabolic equivalents, respectively. Both groups underwent SPU with continuous EMG feedback and completed the interventional oral glucose tolerance test. In the present study, one group completed the interventional oral glucose tolerance test with continuous EMG feedback, while the other group did so without EMG feedback. The duration of the oral glucose tolerance test was 180 minutes in the US study and 120 minutes in the present study.

As with the study conducted by Hamilton et al., our findings indicated that SPU had a beneficial impact on glucose tolerance between minutes 30 and 120 of the oral glucose tolerance test, resulting in a reduction of the area under the curve of blood glucose values during the OGTT by one-third. Furthermore, the beneficial impact of SPU was evident even in the absence of EMG feedback.

In addition to the mechanistic discussion of the metabolic background of SPU by Hamilton et al. [5], the current authors would like to propose a new theory regarding the first-line relevance of the activation of the PIEZO system. This theory suggests that the soleus muscle is not solely a muscle, but rather a proprioceptive organ with high muscle spindle content. This hypothesis is supported by earlier animal studies [13].

This function may be of even greater importance in humans, given the transition from a quadrupedal to a bipedal stance that resulted in a higher need for the contribution of PIEZO2 ion channels to proprioception [14].

Ardem Patapoutian and his team demonstrated that PIEZO2 is the principal mechanosensitive ion channel involved in proprioception [15]. Indeed, mechanotransduction is hierarchical [16], and it is probable that PIEZO2 is the initial activating mechanism in proprioceptive terminals at the onset of exercise, due to its ability to induce a burst of activation [17]. It is also important to consider the possibility of a crosstalk between somatosensory PIEZO2 and peripheral PIEZO1 [18-20]. PIEZO1 has been demonstrated to possess mechanosensory properties in capillaries, and it is postulated that it may play a regulatory role in the distribution of local blood flow through the nervous system [21]. Furthermore, PIEZO1 channels are capable of sensing and responding not only in a spatially restricted manner [22], but also at the whole-body level [23]. This enables them to enhance performance and reset homeostasis, which is regulated by the nervous system through the aforementioned PIEZO2-PIEZO1 crosstalk [24].

The latest research has revealed that PIEZO2 may induce an intrinsic oscillatory interoceptive mechanism via pressure pulse transduction detection, which alters olfactory bulb activity in arousal and is synchronized to brain activities [25]. The hypothesis of long-range synchronization initiated by PIEZO2 was previously postulated in the context of the brain [26] and proposed as a potential mechanism from the periphery, specifically from the muscle spindle to the brain [27, 28]. The new research findings demonstrate the importance of PIEZO2 in arterial pressure pulse detection and provide evidence for an ultrafast oscillatory synchronization mechanism that may be impaired in delayed onset muscle soreness (DOMS).

The relationship between DOMS and amyotrophic lateral sclerosis (ALS) pathomechanism theory is discussed in the aforementioned paper. Accordingly, the transient or irreversible microdamage of PIEZO2 in intrafusal proprioceptive terminals is postulated to be the “gateway to pathophysiology” in DOMS and ALS, respectively [29, 30]. The prominent role of the soleus is also reflected in the induction of the so-called “gateway reflex,” which has the potential to be elicited in other spinal segments, not only in the soleus-innervating L5 segment [31]. It is proposed that the acute transient form of this reflex is the PIEZO2 channelopathy-induced inflammatory reflex [32], a neural circuit-based defense system that maintains neuroimmune homeostasis. In contrast, the chronic form is the aforementioned “gateway reflex,” which involves the “gateway to pathophysiology” in its initial stages [32].

It can therefore be hypothesized that this PIEZO2 channelopathy induces impairment or loss of the long-range proton-based ultrafast synchronization to the hippocampus, both temporarily in the case of DOMS and permanently in ALS [28, 33]. Moreover, this theory of ultrafast proton-based long-range neurotransmission has also been proposed to have a role in first-line insulin regulation [28]. It is also worth noting that this proton-based ultrafast signaling is implicated in the process, with the involvement of catalytically inactive carbonic anhydrase (CA) acting as proton collecting and distributing antennas, as demonstrated in proprioceptive sensory neurons and motoneurons [33]. Noteworthy that animal research indicates that during puberty, the soleus muscle undergoes a transformation from fast-twitch oxidative-glycolytic to slow-twitch oxidative fibers, accompanied by a shift in the status of CA from sensitive to resistant form [34]. The resistant isoform of CA, CA III, is present in the soleus muscle, axons, intrafusal fibers and capillary endothelial cells [34]. This transformation, as postulated by the current authors, is thought to play a role in proton-based ultrafast proprioceptive signaling, given that it acts as a proton-collecting and -distributing antenna. The soleus muscle is a key player in supporting proprioception, postural control and bipedality.

It seems reasonable to suggest that SPU may be an effective exercise method for initiating the PIEZO system in sedentary individuals, by promoting proton-based long-range ultrafast synchronization. It is hypothesized that this ultrafast signaling is impaired in prediabetes. SPU may therefore be an exercise technique to promote this ultrafast proton-based synchronization pathways, thereby leading to more optimal instant insulin regulation in response to mechanotransduction.

Furthermore, unaccustomed or strenuous eccentric contractions have been demonstrated to cause DOMS with associated neuromuscular changes that persist for several days [35-37]. Additionally, the severity of eccentric exercise-induced muscle damage has been shown to correlate with the degree of insulin resistance [38]. It is important to note that DOMS is proposed to be a biphasic non-contact injury mechanism, whereby the primary damage is suggested to be a PIEZO2 channelopathy at intrafusal proprioceptive terminals [30]. Sonkodi et al. attributed the negative correlation to an atypical hippocampal-like metabotropic phospholipase D (PLD)-mGluR on proprioceptive primary afferents, which were first identified by Thomson et al. [39] and are homomeric to metabotropic GluK2 [40]. Indeed, Gluk2 plays a role in the regulation of glucose homeostasis [41]. However, this negative homeostatic regulation is hypothesized to be disrupted by the PIEZO2 channelopathy-induced DOMS effect. Consequently, the impaired PIEZO2-PIEZO1 crosstalk resulting from PIEZO2 channelopathy may lead to the abrogation of proper PIEZO1 modulation, which could in turn result in insulin resistance, given that PIEZO1 is involved in glucose-induced insulin secretion [42]. In support a recent pilot study demonstrated that DOMS inducing exercise protocol impaired orthostasis in a similar fashion like in diabetes [40].

It has been demonstrated that eccentric exercise-induced muscle damage may result in further metabolic alterations. Specifically, it has been observed that the magnitude of the muscle damage correlates with an increase in lipid markers [38]. This phenomenon may be translated as follows: the activation and excitation of PIEZO ion channels may result in lipid and cholesterol depletion in the vicinity of these activated ion channels. The distinctive propeller blade of the PIEZO2 structure is in fact closely surrounded by negatively charged membrane lipids [43-45], and protein-lipid interaction may undergo conformational alterations under allostatic stress [33].

In conclusion, our SPU protocol, which is like that described by Hamilton et al. [5], did not result in the induction of DOMS. Therefore, this exercise regimen appears to facilitate the unidentified PIEZO2-mediated ultrafast proton-based synchronization pathways, thereby promoting more optimal insulin and lipid regulation in response to mechanotransduction, even in individuals with prediabetes.

## Conclusions

1. The physiological effects of SPU can be summarized as follows:
1.1 Blood glucose control
1.1.1 Enhancing glucose uptake: the muscle fibers of the soleus muscle type I contain a substantial number of mitochondria, which are highly efficient in glucose uptake. When the soleus muscle is active, it increases the uptake of glucose from the blood, thereby contributing to a reduction in blood glucose levels.
1. 1.2 Augmentation of insulin sensitivity: regular exercise SPU has been demonstrated to enhance insulin sensitivity, particularly in individuals with insulin resistance. The increase in muscle activity that occurs because of exercise enhances the efficiency of insulin receptors, thereby facilitating the delivery of glucose to muscle cells in a more efficient manner.
1.2. It is recommended that blood circulation be increased.
1.2.1 Venous return: the soleus muscle is often referred to as the “second heart” due to its location in the lower limbs and its function of pumping blood back to the heart. During SPU exercise, the pumping action of the calf muscle stimulates venous return, which improves circulation and reduces venous stasis. This can be of particular importance for individuals who lead sedentary lifestyles or who sit for extended periods of time.
1.3 Enhancing Peripheral Circulation: SPU has been demonstrated to facilitate enhanced circulation to the lower extremities, thereby reducing the incidence of fatigue and edema.
1.3 Enhancing energy production
1.3.1 Mitochondrial activity: the soleus muscle contains a substantial number of mitochondria, which play a pivotal role in the production of energy (ATP). SPU exercise has been demonstrated to enhance mitochondrial activity within the muscle, thereby increasing energy production.
1.3.2 Oxidative Metabolism: Given that the soleus muscle is predominantly composed of slow-contracting muscle fibers, it primarily utilizes oxidative metabolism to meet its energy requirements. This process ensures the production of energy in a more efficient and sustainable manner.
1.4 Increasing endurance and muscle strength
1.4.1 Improving low-intensity endurance: the activation of the soleus muscle has been demonstrated to enhance endurance during low-intensity activities, such as prolonged standing or walking. Although SPU is not as intense as other forms of exercise, the continued activation of the soleus muscle can contribute to improved muscle endurance.
1.4.2 Improving muscle tone: The regular activation of the soleus muscle has been demonstrated to improve muscle tone, which in turn enhances leg strength and stability.
1.5 Enhancing metabolic processes.
1.5.1 Enhance fat metabolism: augmented soleus muscle activity also elevates the oxidation of fatty acids. This can assist in the reduction of body fat and the improvement of lipid profiles (for example, a reduction in LDL cholesterol and an increase in HDL cholesterol).
1.5.2 Increase metabolic rate: it has been demonstrated that regular exercise of SPU may assist in increasing the metabolic rate, particularly in those who lead a sedentary lifestyle.
1.6 A reduction in the inflammatory response is another potential benefit of low-intensity activity. Continuous muscle work, such as that provided by the SPU, has been shown to reduce chronic inflammatory responses. This may be particularly advantageous for individuals with chronic illnesses such as prediabetes or type 2 diabetes.
1.6.2 Reduction of inflammatory markers: Some studies have demonstrated that low-intensity exercise, such as SPU, can assist in the reduction of inflammatory markers (e.g. C-reactive protein, CRP).
1.7 Decrease in venous congestion and swelling: prolonged periods of standing or sitting can result in venous congestion and ankle swelling. The pumping action of the soleus muscle reduces venous stasis, thereby facilitating the drainage of fluids from the legs, which can relieve swelling and pain.
1.8 Summary: The physiological benefits of SPU exercise are numerous and significant. The most notable of these are the control of blood glucose levels, improvements in circulation, increased energy production and enhanced metabolic processes. Such benefits may be particularly advantageous for individuals with prediabetes, insulin resistance, or who spend extended periods sitting. Due to its low intensity, SPU is easily integrated into a daily routine and is a sustainable long-term exercise option.

The objective was to ascertain whether SPU can be an effective intervention for improving glucose regulation in individuals with prediabetes. Furthermore, we examined the feasibility of incorporating SPU into everyday life in the absence of continuous EMG feedback for one group. Furthermore, a potential explanation for this beneficial phenomenon was provided in relation to the PIEZO system.

In conclusion, the SPU exercise can be performed for a minimum of two hours without the onset of fatigue. The performance of SPU has been demonstrated to exert a beneficial effect on blood glucose levels in individuals with prediabetes, even in the absence of EMG feedback. It may be beneficial in the future to test the effectiveness of SPU on a larger sample size without the use of EMG feedback. SPU is a promising form of exercise that can be performed in any location, does not require any equipment, does not cause muscle fatigue even after prolonged performance, is accessible to all, has a systemic effect by activating local contractile elements and is effective in lowering blood glucose levels.

### Limitations of the study

Considering the constraints inherent to our study, it is imperative to underscore the following considerations. It is important to consider the limitations of our study, which involved a relatively small sample size (n=10). No additional demographic data beyond age and sex were available for analysis to ascertain potential associations. The precise execution and application of the SPU movement has only been described in one previous study, therefore the only basis for its education to participants is that single study. The dietary and lifestyle habits of the participants in the days preceding the oral glucose tolerance test were not subject to control. The glucose regulation of the participants was only evaluated following the ingestion of a substantial quantity of exogenous glucose.

## RESOURCE AVAILABILITY

### Lead contact

Further information and requests for resources and reagents should be directed to and will be fulfilled by the lead contact, Dr. Csaba Melczer

## Materials availability

This study did not generate new unique reagents.

## ACKNOWLEDGMENTS

*Financial support for this research was provided by the following sources:* A TKP-2021-EGA-10; A TKP 2021-EGA-37; 2020-1.1.2-PIACI-KFI-2021-00245.

## AUTHOR CONTRIBUTIONS

Conceptualization, Cs.M.; Methodology, D.E., Cs.M.; Investigation, D.E.; Resources, D.E.; Formal Analysis, P.Á.; Writing – Original Draft, D.E., Cs.M.; Writing – Review & Editing, M.T., B.S., G.L.K., Visualization, D.E.; Supervison, B.S.; Project Administration, Cs.M.; Funding Acquisition, Cs.M.;

## DECLARATION OF GENERATIVE AI AND AI-ASSISTED TECHNOLOGIES

## SUPPLEMENTAL INFORMATION

## Box 1. For text boxes, titles are optional

NOTE: (Journal article) and similar text is solely to identify the reference type in each example below; this is not a part of the actual reference format.

## STAR★METHODS

### KEY RESOURCES TABLE

The items in the key resources table (KRT) must also be reported alongside the description of their use in the method details section. Literature cited within the KRT must be included in the references list. Please **do not edit the headings or add custom headings or subheadings** to the KRT. We highly recommend using RRIDs (see https://scicrunch.org/resources) as the identifier for antibodies and model organisms in the KRT. To create the KRT, please use the template below or the <u>KRT webform</u>. See the more detailed <u>Word table template</u> document for examples of how to list items.

| REAGENT OR RESOURCE | SOURCE | IDENTIFIER |
| --- | --- | --- |
| Antibodies |  |  |
| Bacterial and virus strains |  |  |
| Biological samples |  |  |
| Chemicals, peptides, and recombinant proteins |  |  |
| Critical commercial assays |  |  |
| Deposited data |  |  |
| Experimental models: Cell lines |  |  |
| Experimental models: Organisms/strains |  |  |
| Oligonucleotides |  |  |
| Recombinant DNA |  |  |
| Software and algorithms |  |  |
| Other |  |  |

### EXPERIMENTAL MODEL AND STUDY PARTICIPANT DETAILS

Following the provision of initial verbal and subsequent written information, the participant was able to formalize their intention to participate by completing a consent form in accordance with the requirements set out in Decree 23/2002 (9 May 2002) issued by the Ministry of Health and in compliance with the World Medical Association Declaration of Helsinki. The research was approved by the Institutional Research Ethics Committee of the Szent György University Teaching Hospital of Fejér County and the Director General. The study was approved by the National Public Health Center of Hungary (reference number 26914-5/2021/EÜIG).

### METHOD DETAILS

The data collection and interventions for the study were conducted at the Musculoskeletal Rehabilitation Institute of the Szent György University Teaching Hospital of Fejér County in Csákvár during the second half of 2023.

The study’s target population consisted of individuals with prediabetes. Elevated fasting blood glucose and impaired glucose tolerance which are precursors to type 2 diabetes, are separately defined as prediabetic. In accordance with the World Health Organization (WHO) and the Ministry of National Resources’ concurrent delineation of the professional protocol, the criteria for impaired fasting glucose are fasting blood glucose levels between 6.1 and 6.9 mmol/l and blood glucose levels below 7.8 mmol/l at minute 120 of the oral glucose tolerance test. The criteria for IGT are fasting blood glucose levels below 7 mmol/l and between 7.8 mmol/l and 11.1 mmol/l at 120 minutes of the oral glucose tolerance test [6, 7].

The inclusion criteria were as follows: fasting glucose levels between 6.1 and 6.9 mmol/l and blood glucose levels up to 11.1 mmol/l at 120 minutes of the oral glucose tolerance test.

If a participant’s fasting blood glucose level exceeded 7 mmol/l, the OGTT 120 min was equal to or greater than 11.1 mmol/l, and the participant was utilizing glucose tolerance medication or failed to attend any of the scheduled measurements, the participant was excluded from further participation.

The sampling site at the SYNLAB Hungary Limited Liability Company Institute was instrumental in the recruitment of participants and the conduct of control measurements. Individuals who underwent glucose load testing, as indicated by OENO codes 21310 and 23130 at the institute laboratory, were eligible to apply for participation in the study.

A total of 13 individuals expressed their willingness to participate in the trial. The participants were randomly divided into two groups using a software program (Microsoft Corporation, 2402 build version 16.0.17328.20124 64-bit). Seven individuals participated in the control measurement for the group that did not utilize EMG feedback, while five individuals were present for the follow-up measurement. In the group utilizing EMG feedback, six individuals participated in the control measurement and five individuals were present for the follow-up measurement. Following the validation of the exclusion criteria, the measurement results of a total of 10 participants were recorded and subsequently analyzed. The control measurement was identical between the two groups. The oral glucose tolerance test was conducted in the morning, on an empty stomach, following a minimum 10-hour fast. For a period of three days preceding the glucose load, participants were obliged to consume a diet comprising a minimum of 150 grams of carbohydrates per day, with an average level of physical activity. Prior to the commencement of the test, the participants’ blood glucose levels were assessed at a single time point. Should the level fall below 7 mmol/L, the study could proceed. Following the recording of the 0-minute blood glucose values, the subjects were instructed to consume 75 grams of glucose (Magilab Ltd., 1061 Budapest, Király utca 12, CAS number: 50-99-7) dissolved in 250-350 milliliters of water over a five-minute period. During the control measurement, samples were collected at nine time points, with intervals of fifteen minutes for a total of 120 minutes following the initial recording of the 0-minute value. When feasible, blood glucose levels were determined from capillary blood obtained from a warmed hand by pricking the side of the participant’s fingertip. The methodology employed for sampling and the reliability of capillary blood sampling for the detection of blood glucose levels have been validated on an international scale [8, 9]. The procedure was conducted with the use of an Accu-Check Active (Roche Hungary Ltd., Budapest, Hungary) blood glucose monitoring device and test strip, which are in compliance with the current International Organization for Standardization 15197:2013 standard [10]. Among the participants in the group performing SPU without EMG feedback, the 105- and 120-minute sampling results of the control glucose load for one subject were not recorded. At the conclusion of the glucose load, the subject was requested to schedule the subsequent measurement, ideally within a week. At this juncture, a wireless electromyogram electrode (FREEMG 1000, BTS S.p.A., MI, Italy) was positioned on the soleus muscle venter of the lower limb on the side of the subject’s choosing (the muscle venter of the soleus muscle). The five subjects in the group that performed SPU with EMG feedback presented themselves at the study site the morning after the calibration day, after fasting for 10 hours and self-reporting a meal of at least 150 g carbohydrate per day for 3 days prior. We would like to respectfully propose that the participants’ blood glucose levels be measured at the fingertip. If the value was below 6.9 mmol/liter, a wireless EMG electrode was gently placed on the m. soleus muscle venter on the subject’s preferred lower limb, in line with the SENIAM recommendations [11]. The participant was invited to assume a comfortable position in the chair, as had been practiced the previous day. It was suggested that the perpendicular line from the knee joint to the ground should fall on the metatarsophalangeal joint line of the respective side [5, 12]. The participant was given a few minutes to work towards reaching the soleus activity level that had been determined during the preliminary treadmill measurement. This was done by monitoring EMG feedback, which was set to a real-time software frequency of 1000 Hz on a laptop display placed in front of him. This entailed the participant undertaking a plantarflexion movement of approximately 0-45° at a rate of around 60 repetitions per minute. Subsequently, the subject was asked to consume 75 grams of a glucose product dissolved in 250 to 350 milliliters of water over a period of five minutes. Following the ingestion of the sugar solution, we proceeded to take a 0-minute blood glucose measurement and then asked the subject to commence the SPU. The group was provided with real-time EMG feedback throughout the OGTT, which offered continuous insight into soleus muscle activity levels during SPU. Blood glucose measurements were taken at 15-minute intervals over the two-hour oral glucose tolerance test conducted during SPU with EMG feedback. It was encouraged that participants refrain from interrupting the test with any other physically demanding activity if possible. It seems that one participant’s 60th minute blood glucose value was not recorded during the trial. We kindly requested that participants in the trial let us know if they experienced any discomfort, fatigue or muscle cramps. We are pleased to report that no complaints were made during the trial that would have required us to suspend the trial. The group that performed SPU without EMG feedback followed the same preparation and blood glucose measurement as the other group, followed by OGTT during the intervention (SPU), but unfortunately, no real-time EMG feedback was available for this group. It was necessary for them to estimate the level of soleus activity and range of motion of the movement, which had previously been practiced for only a few minutes and measured during gait. This was based on their experience up to that point. It would have been beneficial to correct the SPU execution during the OGTT period. We would like to kindly remind you that blood glucose measurements were taken every 15 minutes for the duration of the two-hour oral glucose tolerance test. It was kindly requested that participants refrain from engaging in any other physically demanding activities during the test, if possible. Unfortunately, there was a technical issue during the trial which meant that one participant’s results were not recorded from the 75th minute of the test. We kindly requested that participants report any discomfort, fatigue or muscle cramps immediately. We are grateful that during the study, subjects did not report any complaints that would have required us to suspend the study. The flowchart of the study design is shown in Figure 4.

**Figure 4.**
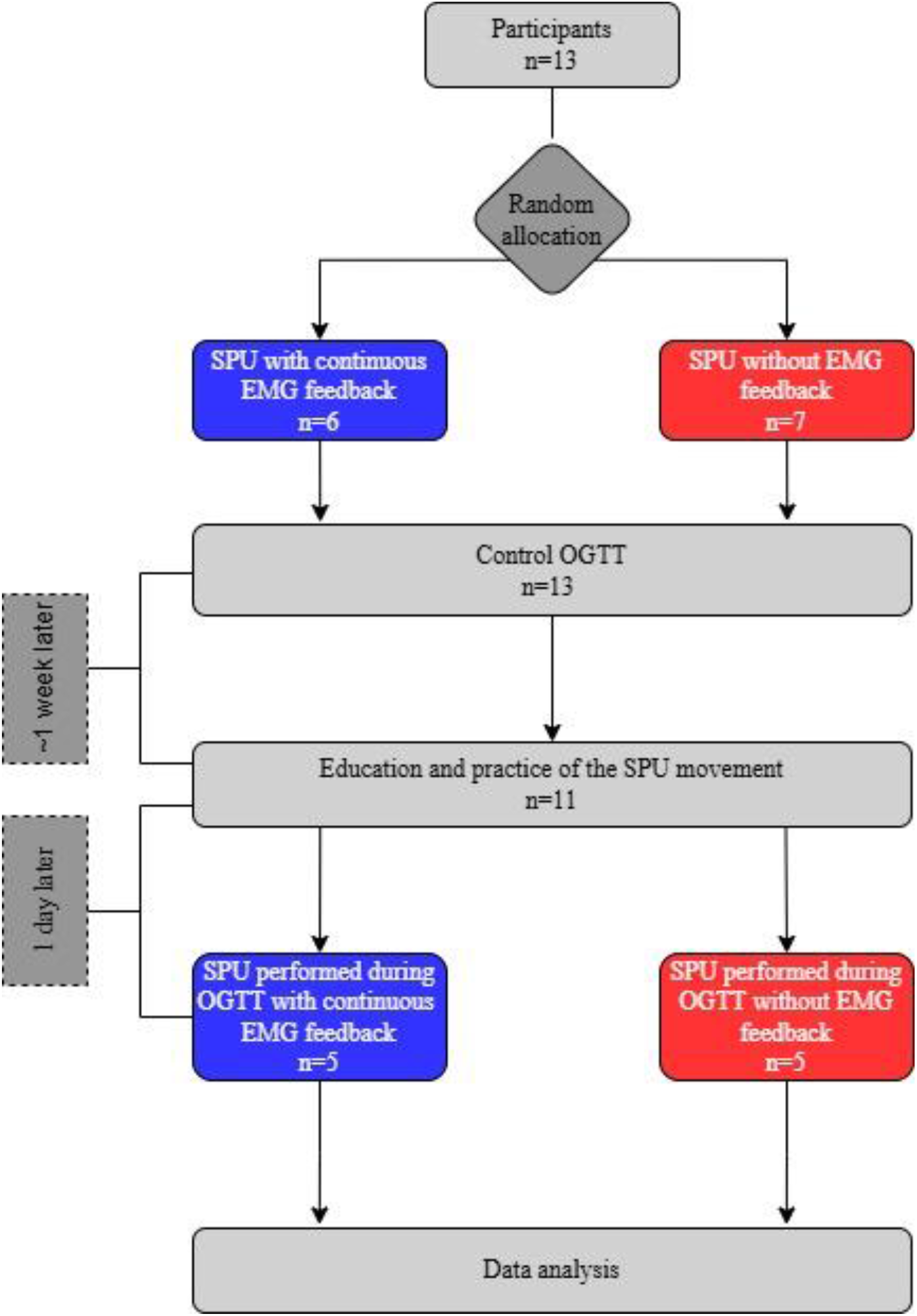
Study design.

### QUANTIFICATION AND STATISTICAL ANALYSIS

For the statistical data analysis, we used Microsoft Excel (Microsoft Corporation, 2402 build version 16.0.17328.20124 64-bit) to record the data and perform simple mathematical operations. In order to determine the area under the curve defined by the blood glucose levels measured every 15 minutes during the oral glucose tolerance tests, as well as to test the statistical significance of the degree of change in blood glucose levels at a given measurement time point of the oral glucose tolerance test within group and between the two groups, a mixed effects model with Tukey’s multiple comparison test at the 95% confidence level were used. The analysis was conducted using GraphPad Prism 10 (GraphPad Software, Boston, Massachusetts USA). To examine the change in area under the curve within and between groups, we employed a one-factor analysis of variance at the 95% confidence level, utilizing the statistical software SPSS (IBM Corp. Released 2019. IBM SPSS Statistics for Windows, Version 26.0. Armonk, NY: IBM Corp.).

### ADDITIONAL RESOURCES

